# A cross taxonomic comparison of bird and butterfly communities of Tamhini Wildlife Sanctuary across two decades

**DOI:** 10.1101/2022.01.30.478416

**Authors:** Shawn Dsouza, Anand Padhye

## Abstract

Human disturbance can alter the structure and function of ecological communities. We studied bird and butterfly communities of Tamhini wildlife sanctuary to understand the effects two decades of changing land use and management. We replicated a previous study conducted between 1998 and 2001, sampling 7 line transects every fortnight between April 2016 and April 2017. We analyzed the diversity and structure of functional groups and compared our findings to previous surveys. Species diversity increased for both taxa, however, community composition was significantly different across studies. Changing land may affect multiple trophic levels and alter community assembly and ecosystem function.

## Introduction

Habitat modification and disturbance are major threats to biodiversity in the tropics. Large swaths of land have been converted for monoculture plantations and unsustainable logging practices threaten endemic species in biodiverse regions ^1,2^. Large areas of pristine habitats that are required for the maintenance of diversity and ecosystem functioning are now hard to come by ^3^. In the face of these threats, it is imperative that our management practices are dynamic and that their efficacy can be monitored continuously.

The structure of ecological communities is determined by the interaction of global as well local processes, namely, speciation, dispersal, selection, and drift ^4^. Human activities such as converting land for agricultural use, logging and infrastructure development alter these processes resulting in the loss of diversity and in the long run, the loss of ecosystem services as well ^2,3^. Due to the complexity of interactions in ecological communities and the multifaceted nature of human activities, it is difficult to quantify the complete effects of human disturbance in pristine ecosystems ^5^.

While monitoring ecological poses a significant challenge, ecological indicators offer an economical solution. Ecological indicators are species that have characteristics (such as sensitivity to pollutants or specific habitat requirements) relevant to monitoring the ecosystem functions of an area ^6^. Their use has risen in the past few decades ^7^. Indicator species can serve as useful indices for the selection of areas to be conserved and for the effective allocation of management resources ^8^. They can also serve as feedback for adaptive management of protected areas ^9^.

Monitoring multiple indicator taxa offers a more holistic overview of the effects of disturbance ^10^. The efficacy of birds and butterflies as ecological indicators of human disturbance and habitat modification has been well studied (eg. ^1,11–13^). In addition, the effects of disturbance on these taxa differ due to differences in their ecology. Birds have complex feeding habits and respond to changes in habitat structure. Butterflies on the other hand respond to local level changes in parameters such as the composition of vegetation ^14,15^.

There are also considerable differences in the monitoring of community parameters. Measuring changes or differences in diversity can be misleading and responses may vary by taxa. For example, Hill and Hammer (2004) found that butterfly diversity may increase in response to site level disturbance, whereas bird diversity reduces with increasing local disturbance ^14^. Analyzing the response of components of the community such as functional guilds can reveal specific patterns ^13^. For example, insectivorous birds were disproportionately affected due to unsustainable logging ^16^.

The Western Ghats is a biodiversity rich region that is threatened by encroachment through activities such as infrastructure development and agriculture ^17^. In addition, many people in the region also depend on forest resources and non-timber forest products. The Western Ghats provide vital ecosystem services to much of western India ^18,19^. Tamhini is a recently established wildlife sanctuary in the Northern Western Ghats (National Green Tribunal of India, 2015) that has a complex history of protection and management. Despite its protected status the area is also threatened by encroachment and tourist activities.

We aimed to understand the effects of changing land use and management practices across taxa in Tamhini WLS over two decades. In the present study, our objectives were: (a) to compare community shifts (diversity and composition) across indicator taxa at two trophic levels, birds and butterflies, (b) to determine differences in magnitude of shifts across functional groups within these taxa, and (c) to determine the effect of these shifts on community function.

## Materials and Methods

### Study Area

Tamhini Wildlife Sanctuary (18°27’N, 73°25’E) is situated in the Northern Western Ghats around 60 kms west of Pune, Maharashtra (Figure 1). The area is dominated by hilly terrain and an average elevation of 600m asl. A large part of the study area has been modified for human use ranging from, farmlands and pastures to resorts and residential complexes ^20^. The climate is moderate and tropical throughout most of the year, with heavy to torrential rains (5500 – 6500 mm) during the monsoons as is the case with much of the Western Ghats ^21^. Mulshi Lake, situated in the sanctuary is a large reservoir fed by the Mula and Neela rivers, which retains water throughout the year, maintaining both favorable temperatures and humidity in the area.

**Figure 1:**
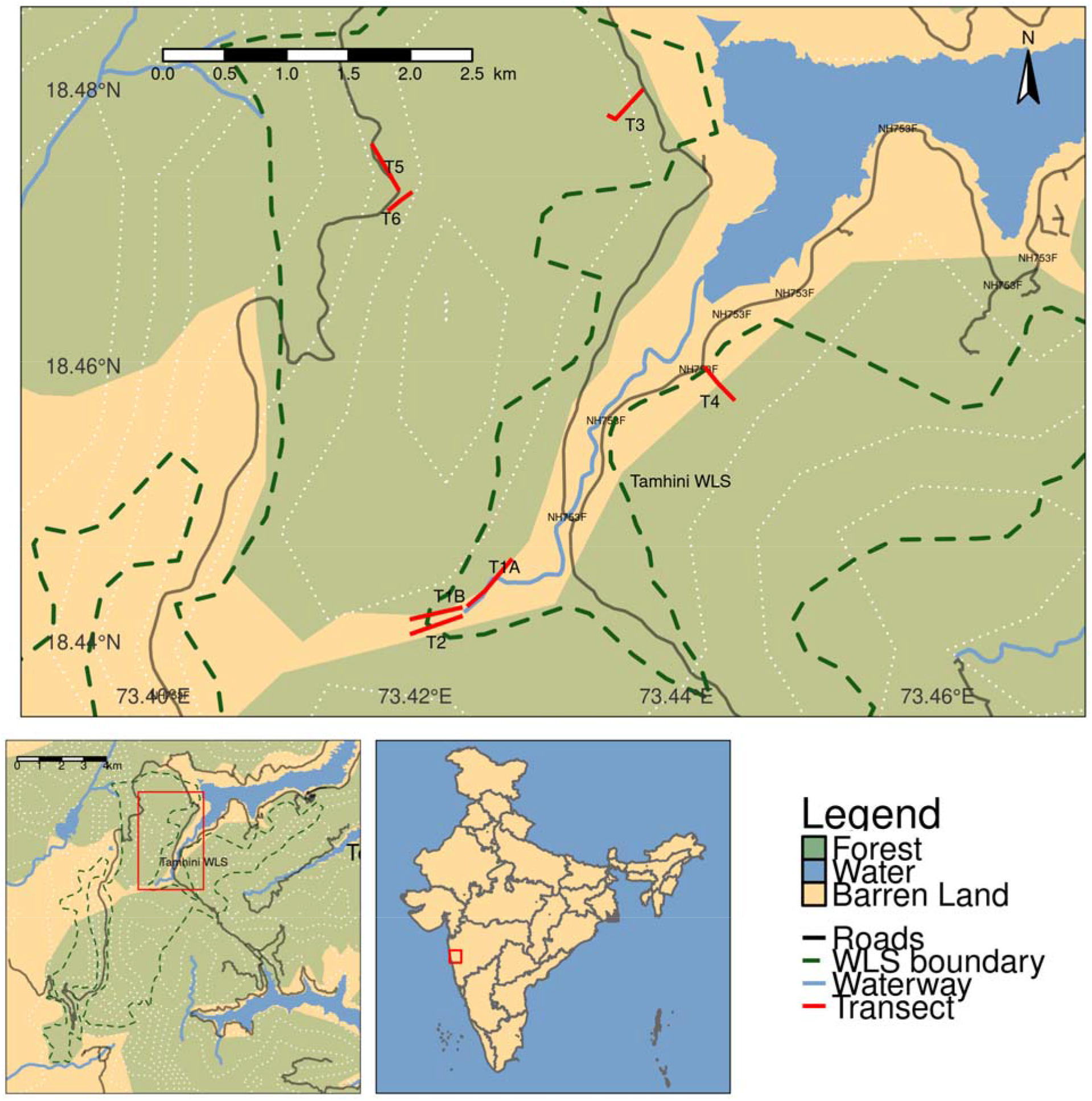
Location of transects (top) laid in Tamhini Wildlife Sanctuary (bottom left), situated around 60k west of Pune in the Northern Western Ghats (bottom center). White dotted lines indicate 100m elevation contours.

### Data Collection and Sampling

We replicated the studies conducted by Padhye et al. between 1998 and 2001 ^22,23^. Seven line transects were laid out throughout the study area (Fig. 1) identical to the previous studies ^22,23^. These represented 4 habitats, namely, 1. Riparian, 2. Evergreen Forest, 3. Human Habitation and Cultivated Land and 4. Scrubland and Grassland (Figure 1). Transects were sampled every fortnight between April 2016 and April 2017, for a total of 24 visits. Bird and adult butterfly abundances were recorded along each of these transects between 7 am – 11 am and 4pm – 6pm, when the subjects are most active and there is a chance to encounter even crepuscular species ^24^. The number of visits and sampling effort were similar between the current and previous studies. Photographs were taken when additional diagnosis was required. Seasonal changes in land use and vegetation were also recorded incidentally.

### Data Analysis

We calculated the diversity of both taxa in each site as effective number of species (D^1^) ^25^. We compared the change in diversity across studies using a linear model with sites as samples. We visualized community composition across studies using Non-metric dimensional scaling ^26^. We then compared change in community composition across studies using a Permutative Analysis of Variance test ^27,28^.

We collected information on host plant species and families for the butterfly species we encountered during our survey as well as from the previous study using the HOSTS database of the Natural History Museum, London ^29^. We then classified butterfly species into trophic guilds based on host plant habit as Grass, Herb, Liana, Shrub, Tree Specialist or Generalist ^13,30^. Similarly, we classified birds into guilds based on diet data from the Birds of the World Database ^31^. The bird species were classified as Carnivores, Frugivores, Granivores, Insectivores, or Omnivores. We compared change in functional diversity across studies using linear model for each trophic guild in each taxon.

We tested for change in community function by first calculating the habitat specialization index (HSI) for each species of both taxa as 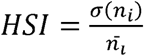, where n_i_ is the relative proportion of each species in each habitat. We also computed a trophic specialization index (TSI) for each species as 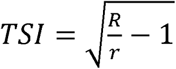, where R is the total number of host plants or prey types used by the community and r is the number of host plants or prey types used by a species. We then calculated a community specialization index as the mean of the individual species specialization indices at each site ^32^. We compared change in community specialization indices across studies using linear models with sites as samples ^33^.

All our analysis was carried out in R version 4.1 (R Core Team 2016). The data collected and/or analyzed during this study as well as the code for analysis is available at https://github.com/cheesesnakes/tamhini-birds-butterflies.

## Results

### Species diversity and composition of birds and butterflies

We encountered 105 bird species (N = 2021) and 66 butterfly species (N = 2014) in 2016 - 2017 compared to 70 bird species (n = 1007) and 45 butterfly species (n = 515) in 1998 – 2001. The species diversity of birds increased significantly compared to 1998 - 2001 (D^1^_1998-2001_ = 17.31 ± 6.47, D^1^_2016-2017_ = 24.88 ± 5.74, β = 7.56 ± 3.38, T_11_ = 2.23, p = 0.04, r^2^ = 0.31). However, the change in butterfly diversity was not significant (D^1^_1998-2001_ = 15.87 ± 3.42, D^1^_2016-2017_ = 20.4 ± 3.24, β = 4.53 ± 2.07, T_8_ = 2.18, p = 0.056, r^2^ = 0.34, Figure 2). The species composition of both taxa also changed significantly over the past two decades (C_birds_ = 0.55, R^2^_birds_ = 0.25, p_birds_ = 0.001, C_butterflies_ = 0.67, R^2^_butterflies_ = 0.25, p_butterflies_ = 0.02, Figure 3).

**Figure 2:**
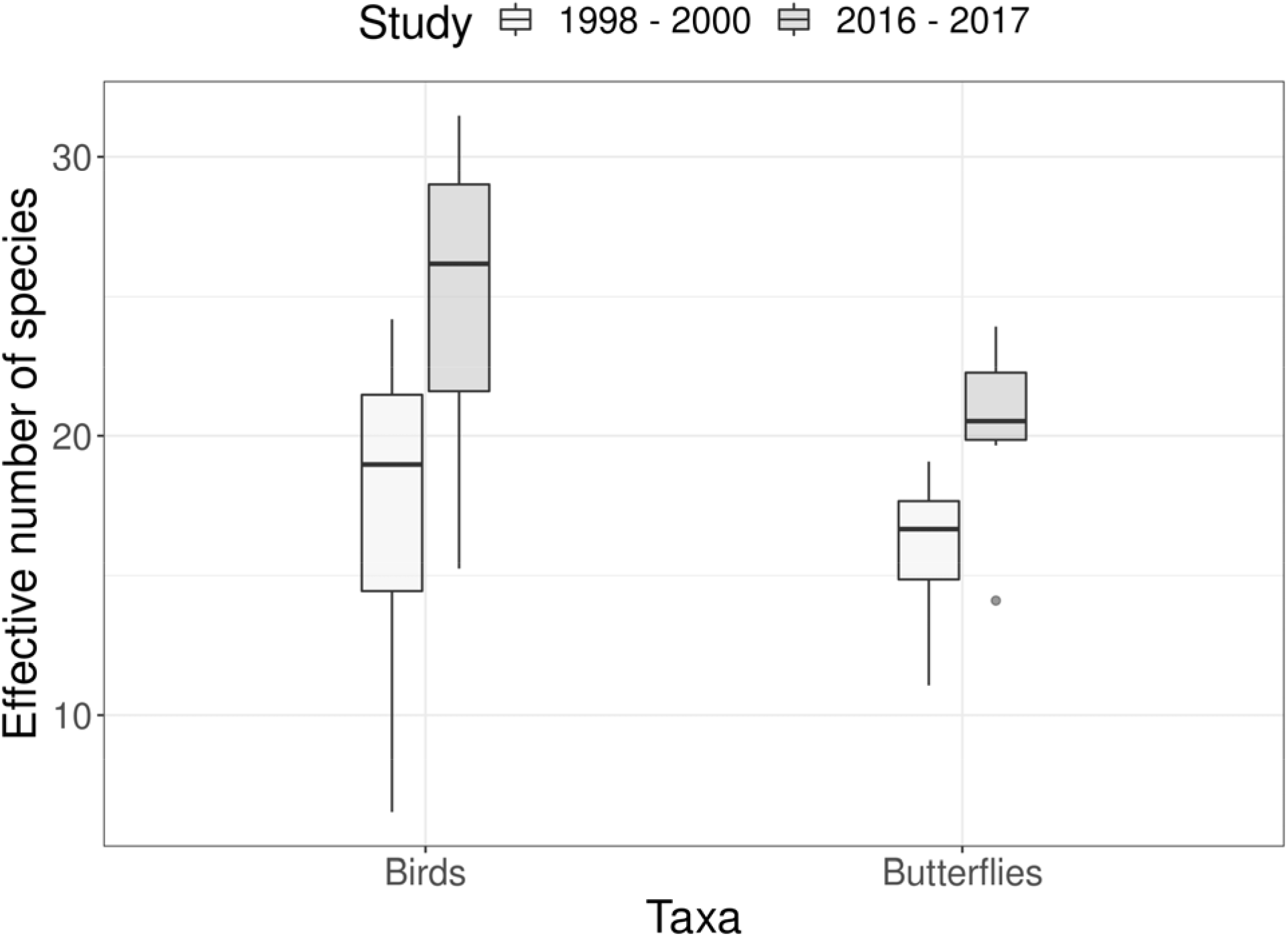
Change in first order diversity of bird and butterflies in Tamhini WLS between 1998 and 2017.

**Figure 3:**
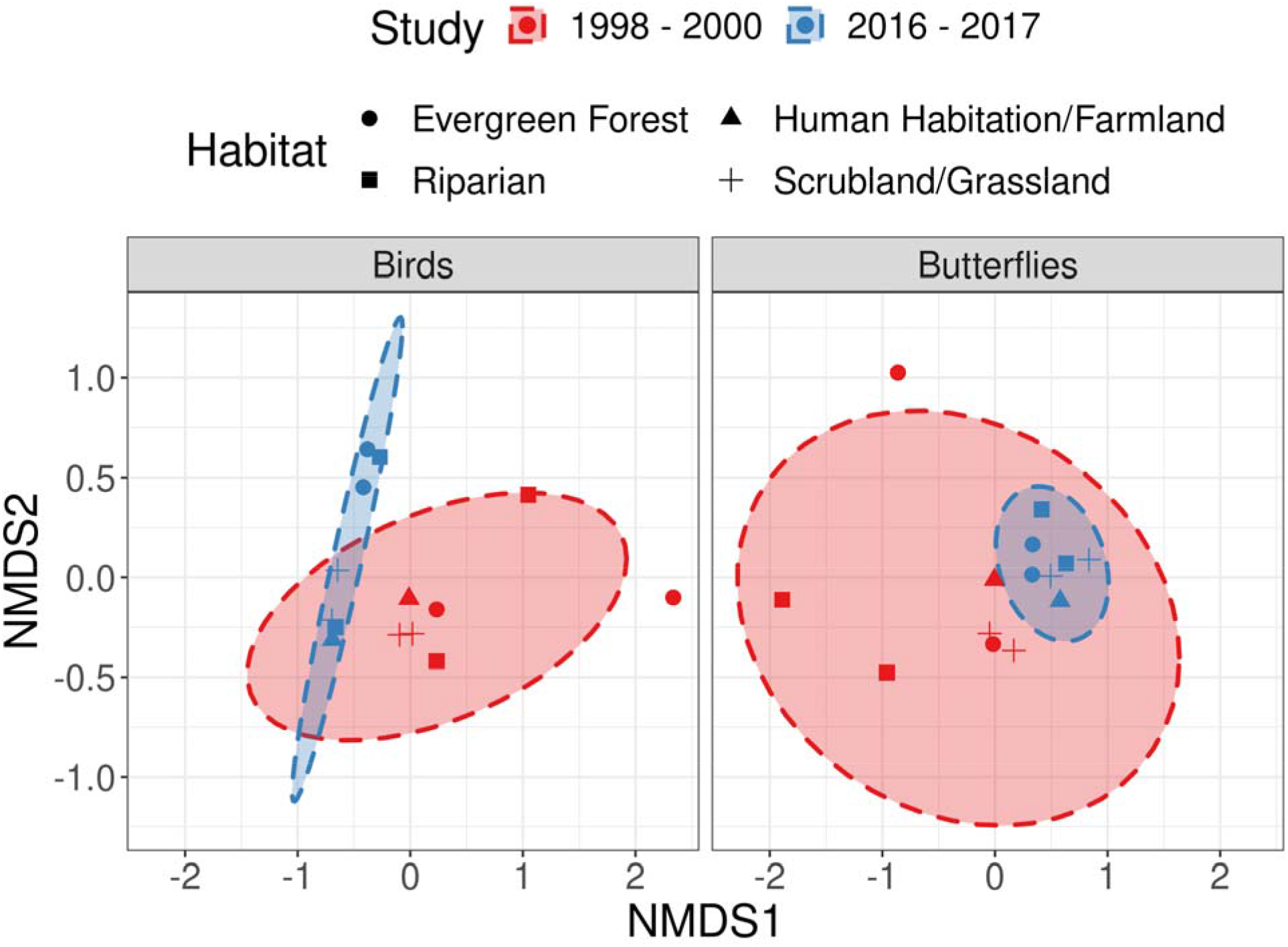
NMDS plot depicting change in community composition of bird and butterflies in Tamhini WLS between 1998 and 2017.

### Functional diversity of birds and butterflies

The diversity of insectivorous, carnivorous, and omnivorous birds increased significantly at Tamhini WLS when compared to the previous study. However, the diversity of granivorous and frugivorous birds was not significantly different. It should be noted that sample sizes for carnivorous birds was low. Insectivorous birds saw the greatest increase in diversity out of all bird trophic guilds.

The diversity of grass specialist and generalist butterflies increased significantly. On the other hand, the diversity of herb specialist, shrub specialist and tree specialist species was not significantly different. Generalist butterflies saw the greatest increase in diversity. Liana specialists were only encountered in the previous study and were not encountered in the current study (Table 1, Figure 4).

**Table 1:**
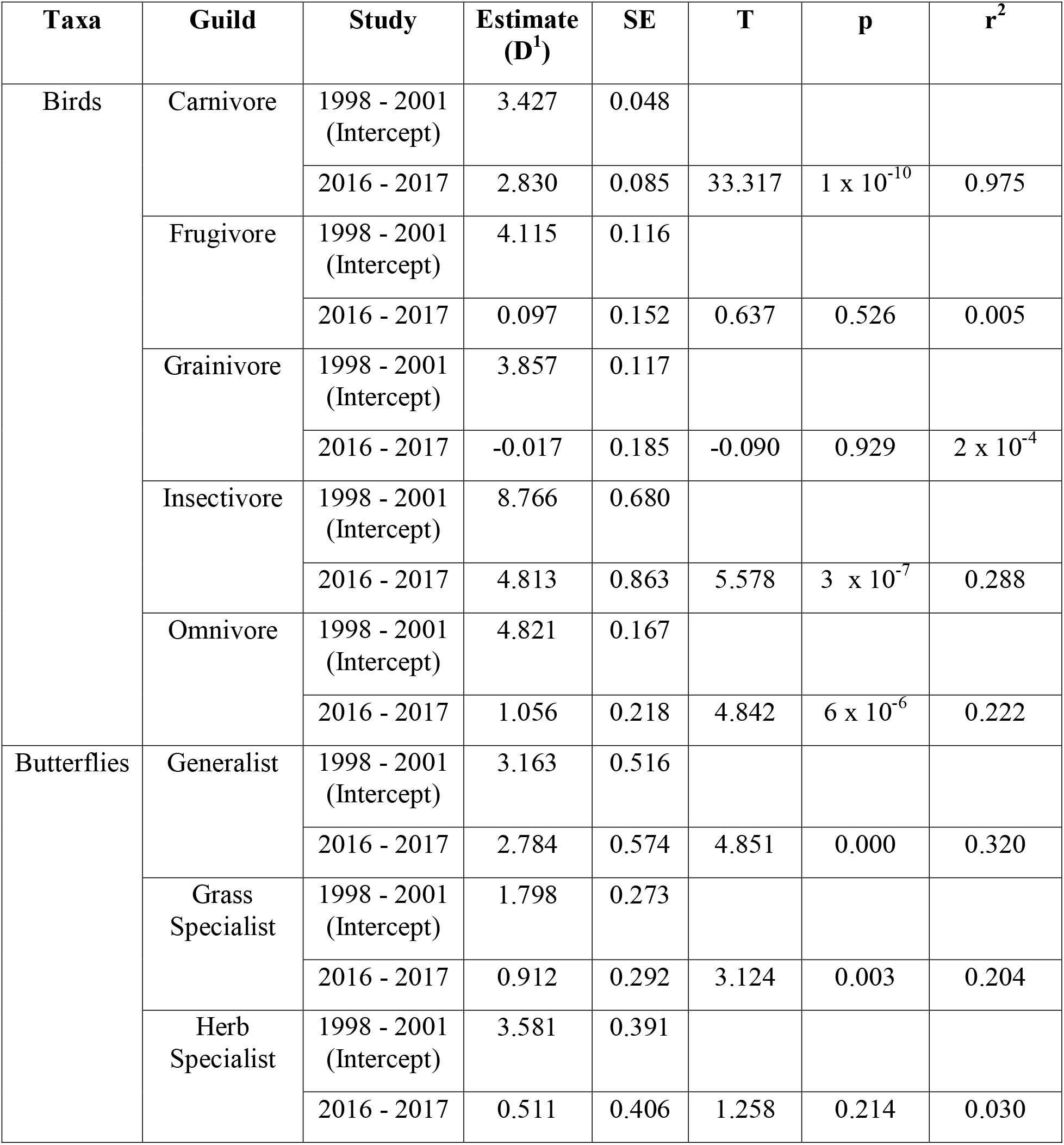
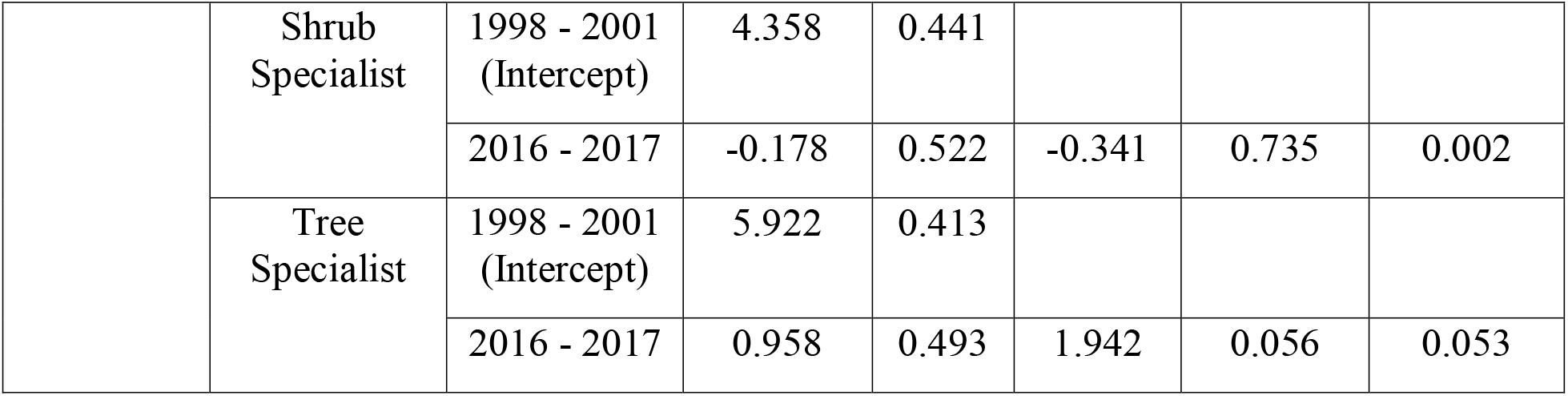
Comparing diversity of trophic guilds in bird and butterfly communities at Tamhini WLS between 1998 and 2017.

**Figure 4:**
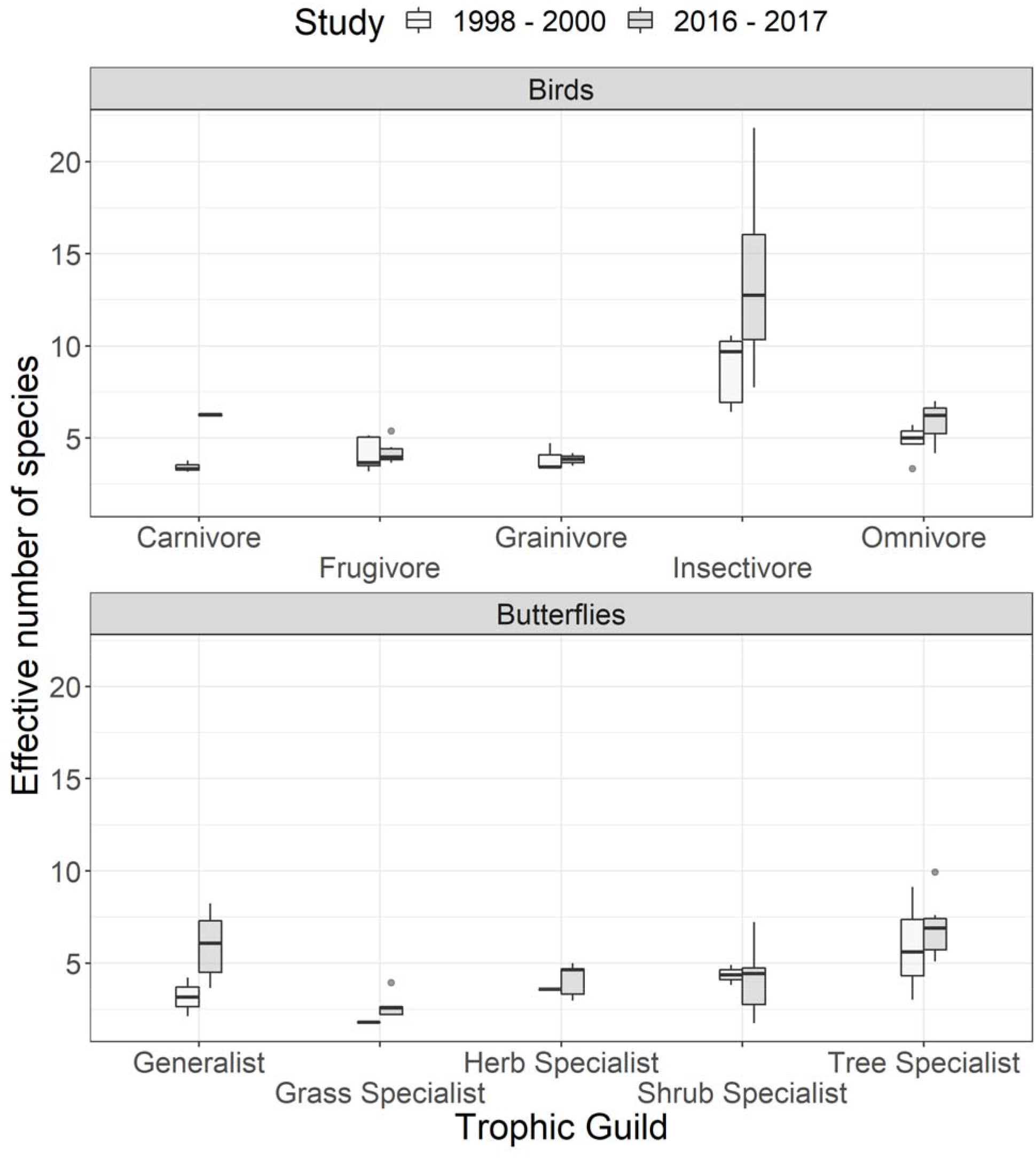
Change in relative proportion trophic guilds of birds and butterflies in Tamhini WLS between 1998 and 2017. Liana specialist butterflies were only encountered in the current study and not the previous study. In addition, our sample of liana specialists was too small to compute diversity metrics, and have thus been excluded from the figure and analysis.

### Effect on community function

Neither birds (CSI^T^_1998-2001_ = 2.58 ± 0.16, CSI^T^_2016-2017_ = 2.58 ± 0.09, β = -0.05 ± 0.06, T_9_ = 0.81, p = 0.43, r^2^ = 0.05) nor butterflies (CSI^T^_1998-2001_ = 5.33 ± 0.39, CSI^T^_2016-2017_ = 5.06 ± 0.24, β = - 0.27 ± 0.17, T_9_ = 1.57, p = 0.14, r^2^ = 0.17) showed a significant change in mean trophic niche width. However, butterflies showed a slight trophic niche contraction. Similarly, the degree of community habitat specialization of both bird (CSI^H^_1998-2001_ = 0.54 ± 0.1, CSI^H^_2016-2017_ = 0.49 ± 0.02, β = -0.042 ± 0.039, T_9_ = 0.81, p = 0.43, r^2^ = 0.08) and butterfly (CSI^H^_1998-2001_ = 0.61 ± 0.09, CSI^H^_2016-2017_ = 0.57 ± 0.03, β = -0.041 ± 0.034, T_9_ = 1.57, p = 0.14, r^2^ = 0.1) communities was slightly lower but not significantly different.

## Discussion

Tamhini WLS is a biodiversity rich area, supporting many bird, butterfly, amphibian and reptile species in a relatively small area. Despite its protected status the area is under threat of encroachment ^22,23,34^. Tamhini also has an interesting history of management interspersed with privately owned land, reserved forest, and human habitations. In addition, local people still depend on the remaining forests for firewood and non – timber forest products (pers. obs.).

Our comparison of bird and butterfly communities over two decades revealed a significant increase in diversity of birds and though not significant, an increase in the diversity of butterflies as well (Figure 2). An increase in diversity does not necessarily mean that management practices are effective. Different taxa can respond differently to disturbance and these effects vary across spatio-temporal scales ^10,33,35^. Both bird and butterfly communities displayed significant turnover when compared across studies (Figure 3). This change in community composition can be attributed to a variety of underlying processes including change in land use patterns, changes in habitat structure, natural cyclic variation, and sensitivity of subsets (e.g., functional groups) of the community ^5^.

When we break down the diversity of these species into functional components, we can observe more fine scale patterns. Trophic and habitat specialization reduced slightly but not significantly for both taxa. In the case of butterfly communities, we observed a large increase in generalist species diversity compared to more specialist species in our study (Table 1, Figure 4). Butterflies are sensitive to disturbance at smaller spatial scales, particularly changes to vegetation ^15^. Species specializing in specific caterpillar host plants may be at a disadvantage in the face of human activities that alter plant communities such as logging and slash and burn agriculture ^13^, both of which were observed on our transects. On the other hand, bird communities in Tamhini saw a large increase in insectivorous birds in addition to a moderate increase in omnivorous bird species (Table 1, Figure 4). Habitat modification such as converting land for agriculture can result in changes in resource availability and consequent changes in interspecific completion ^36^. Thus, species with specific resource requirements may be at an advantage in human modified landscapes^33,37^.

Our study is limited in spatial scale and thus our inferences are difficult to generalize beyond the case of Tamhini WLS. In addition, sampling effort differed among the studies compared. Sampling effort (in terms of number of individuals sampled) can have a large impact on both alpha and beta diversity and must be kept in mind when interpreting results (see supplementary materials). Differences in expertise in identification of the focal taxa may also introduce additional biases. However, the temporal scale of our comparison reveals useful insights into how these communities respond to human presence. Future studies may benefit from better spatial replicates and more even sampling effort to detect effects that we were not able to in the current study.

Despite an apparent increase in diversity at the community level, we observed a shift in functional diversity across both bird and butterfly communities in Tamhini WLS. Such shifts may have implications for community assembly and ecosystem function. Looking beyond species diversity may prove useful for the management of biodiverse areas in the Western Ghats such as Tamhini WLS.

## Acknowledgments

We’d like to thank the Department of Biodiversity, Abasaheb Garware College, Pune for their help and mentorship. We thank Dr. Kartik Shanker for his comments on the manuscript. We extend our appreciation to Tarun Menon, Anand Pendharkar, Rahul Pungaliya and all others who offered their help in the form of ideas and support. We also thank all those who helped SD in data collection including but not limited to Akshay Marathe, Aseem Shendye, Tarun Menon, Srushti Bhave, Vishal Varma and Pranav Mehsalkar.

## Notes

https://github.com/cheesesnakes/tamhini-birds-butterflies

### Competing Interest Statement

The authors have declared no competing interest.

## Literature Cited

1. Hill, J. K., Hamer, K. C., Lace, L. A., Banham, W. M. T., Lacet, L. A., and Banhamt, W. M. T., Effects of Selective Logging on Tropical Forest Butterflies on Buru, Indonesia. Source J. Appl. Ecol. J. Appl. Ecol., 1995, 32, 754–760.

2. Sodhi, N. S., Koh, L. P., Clements, R., et al., Conserving Southeast Asian forest biodiversity in human-modified landscapes. Biol. Conserv., 2010, 143, 2375–2384.

3. Anand, M. O., Krishnaswamy, J., Kumar, A., and Bali, A., Sustaining biodiversity conservation in human-modified landscapes in the Western Ghats: Remnant forests matter. Biol. Conserv., 2010, 143, 2363–2374.

4. Vellend, M., Conceptual Synthesis in Community Ecology. Q. Rev. Biol., 2010, 85, 183– 206.

5. Mittelbach, G. G. and McGill, B. J., Biodiversity and ecosystem functioning Community Ecol., Oxford University Press, 2012.

6. Landres, P. B., Verner, J., Thomas, J. W., Landres, P. B., and Verner, J., Society for Conservation Biology Ecological Uses of Vertebrate Indicator SpeciesLJ: A Critique Ecological Uses of Vertebrate Indicator SpeciesLJ: A Critique. 1988, 2, 316–328.

7. Siddig, A. A. H., Ellison, A. M., Ochs, A., Villar-Leeman, C., and Lau, M. K., How do ecologists select and use indicator species to monitor ecological change? Insights from 14 years of publication in Ecological Indicators. Ecol. Indic., 2016, 60, 223–230.

8. Caro, T. M. and Doherty, G. O., On the Use of Surrogate Species in Conservation Biology. 1999, 13, 805–814.

9. Dufrene, M., Legendre, P., Monographs, S. E., and Aug, N., Species Assemblages and Indicator SpeciesLJ: The Need for a Flexible Asymmetrical Approach SPECIES ASSEMBLAGES AND INDICATOR SPECIESLJ: THE NEED FOR A FLEXIBLE ASYMMETRICAL APPROACH. 2015, 67, 345–366.

10. Lawton, J. H., Bignell, D. E., Bolton, B., et al., Biodiversity inventories, indicator taxa and effects of habitat modification in tropical forest. Nature, 1998, 391, 72–76.

11. Canterbury, G. E., Martin, T. E., Petit, D. R., Petit, L. J., and Bradford, D. F., Bird communities and habitat as ecological indicators of forest condition in regional monitoring. Conserv. Biol., 2000.

12. Blair, R. B., BIRDS AND BUTTERFLIES ALONG AN URBAN GRADIENTLJ: SURROGATE TAXA FOR ASSESSING BIODIVERSITYLJ? 1999, 9, 164–170.

13. Cleary, D. F. R., Cleary, D. F. R., Boyle, T. J. B., et al., The impact of logging on the abundance, species richness and community composition of butter y guilds in Borneo. Jen, 2005, 129, 52–59.

14. Hill, J. K. and Hamer, K. C., Determining impacts of habitat modification on diversity of tropical forest fauna: the importance of spatial scale. J. Appl. Ecol., 2004, 41, 744–754.

15. Debinski, D. M., VanNimwegen, R. E., and Jakubauskas, M. E., Quantifying Relationships Between Bird And Butterfly Community Shifts And Environmental Change. Ecol. Appl., 2006, 16, 380–393.

16. Sreekar, R., Srinivasan, U., Mammides, C., et al., The effect of land-use on the diversity and mass-abundance relationships of understory avian insectivores in Sri Lanka and southern India. Nat. Publ. Group, 2015.

17. Jha, C. S., Dutt, C. B. S., and Bawa, K. S., Deforestation and land use changes in Western Ghats, India. Curr. Sci., 2000, 79, 231–238.

18. Gadgil, M., Documenting Diversity. Curr. Sci., 1996, 70, 36–44.

19. Gadgil, M., Berkes, F., and Folke, C., Indegenous knowledge for biodiversity conservation. Ambio, 1993, 22, 151–156.

20. Padhye, A., Shelke, S., and Dahanukar, N., Distribution and composition of butterfly species along the latitudinal and habitat gradients of the Western Ghats of India. Check List, 2012, 8, 1196–1215.

21. Kothawale, D. R., Despande, N. R., Narkhedkar, S. G., and Kulkarni, J. R., An unidentified heavy rainfall station ‘Tamhini’ in the northern region of Western Ghats of India. Int. J. Climatol., 2017, 37, 1416–1431.

22. Padhye, A. D., Dahanukar, N., Paingankar, M., Deshpande, M., and Deshpande, D., Season and landscape wise distribution of butterflies in Tamhini, northern Western Ghats, India. Zoos Print J., 2006, 21, 2175–2181.

23. Padhye, A. D., Paingankar, M., Dahanukar, N., and Pande, S., Season and landscape element wise changes in the community structure of avifauna of Tamhini, Northern Western Ghats, India. Pap. Zoos Print J., 2007, 22, 2807–2815.

24. Kunte, K. J., Seasonal patterns in butterfly abundance and species diversity in four tropical habitats in northern Western Ghats. J. Biosci., 1997, 22, 593–603.

25. Jost, L., Entropy and diversity. Oikos, 2006, 113, 363–375.

26. Kruskal, J. B., Multidimensional scaling by optimizing goodness of fit to a nonmetric hypothesis. Psychometrika, 1964, 29, 1–27.

27. Oksanen, J., Multivariate analysis of ecological communities in R: vegan tutorial. R Doc., 2015, 43.

28. Anderson, M. J. and Walsh, D. C. I., PERMANOVA, ANOSIM, and the Mantel test in the face of heterogeneous dispersions: What null hypothesis are you testing? Ecol. Monogr., 2013, 83, 557–574.

29. Robinson, G. S., Ackery, P. R., Kitching, I. J., Beccaloni, G. W., and Hernandez, M., HOSTS -A Database of the World’s Lepidopteran Hostplants. Natural History Museum, London. Database, 2010.

30. Janz, N. and Soren, N., Butterflies and plants: a phylogenetic study. Evolution, 1998, 52, 486–502.

31. Billerman, S. M., Keeney, B. K., Rodewald, P. G., and Schulenberg, T. S., Birds of the World. Cornell Laboratory of Ornithology, Ithaca, NY, USA. Database,2020.

32. Julliard, R., Clavel, J., Devictor, V., Jiguet, F., and Couvet, D., Spatial segregation of specialists and generalists in bird communities. Ecol. Lett., 2006, 9, 1237–1244.

33. Devictor, V. and Robert, A., Measuring community responses to large-scale disturbance in conservation biogeography. Divers. Distrib., 2009, 15, 122–130.

34. Dahanukar, N. and Padhye, A., Amphibian diversity and distribution in Tamhini, northern Western Ghats, India. Curr. Sci., 2005, 88, 1496–1501.

35. Bowman, D. M. J. S., Woinarski, J. C. Z., Sands, D. P. A., Wells, A., and McShane, V. J., Slash-and-Burn Agriculture in the Wet Coastal Lowlands of Papua New Guinea: Response of Birds, Butterflies and Reptiles. J. Biogeogr., 1990, 17, 227–239.

36. Maron, M., Main, A., Bowen, M., et al., Relative influence of habitat modification and interspecific competition on woodland bird assemblages in eastern Australia. Emu - Austral Ornithol., 2011, 111, 40–51.

37. Posa, M. R. C. and Sodhi, N. S., Effects of anthropogenic land use on forest birds and butterflies in Subic Bay, Philippines. Biol. Conserv., 2006, 129, 256–270.

