## Supplementary Materials for "A cross taxonomic comparison of bird and butterfly communities of Tamhini Wildlife Sanctuary across two decades"

Table A: Number of birds and butterflies (individuals) sampled in 1998 – 2000 and 2016 – 2017. (T = Transect)

| Study | Taxa | T1A | T1B | T2 | T3 | T4 | T5 | T6 |
| --- | --- | --- | --- | --- | --- | --- | --- | --- |
| 1998 - 2000 | Birds | 255 | 165 | 103 | 213 | 236 | 7 | 28 |
| 1998 - 2000 | Butterflies | 93 | 59 | 12 | 180 | 157 | 10 | 4 |
| 2016 - 2017 | Birds | 480 | 393 | 358 | 289 | 165 | 203 | 133 |
| 2016 - 2017 | Butterflies | 432 | 188 | 207 | 652 | 187 | 224 | 124 |

Table B: List of sites sampled with habitat attributes.

| Site | Habitat | Start Latitude | Start Longitude | End Latitude | End Longitude |
| --- | --- | --- | --- | --- | --- |
| T1A | Human Habitation/Farmland | 18.446017 | 73.427433 | 18.442553 | 73.423978 |
| T1B | Scrubland/Grassland | 18.442493 | 73.423636 | 18.441612 | 73.419593 |
| T2 | Riparian | 18.440535 | 73.419611 | 18.441869 | 73.42365 |
| T3 | Scrubland/Grassland | 18.48002 | 73.437459 | 18.477847 | 73.435341 |
| T4 | Evergreen Forest | 18.459945 | 73.442139 | 18.457477 | 73.444475 |
| T5 | Evergreen Forest | 18.472681 | 73.418793 | 18.476085 | 73.416667 |
| T6 | Riparian | 18.471213 | 73.417911 | 18.472598 | 73.419794 |

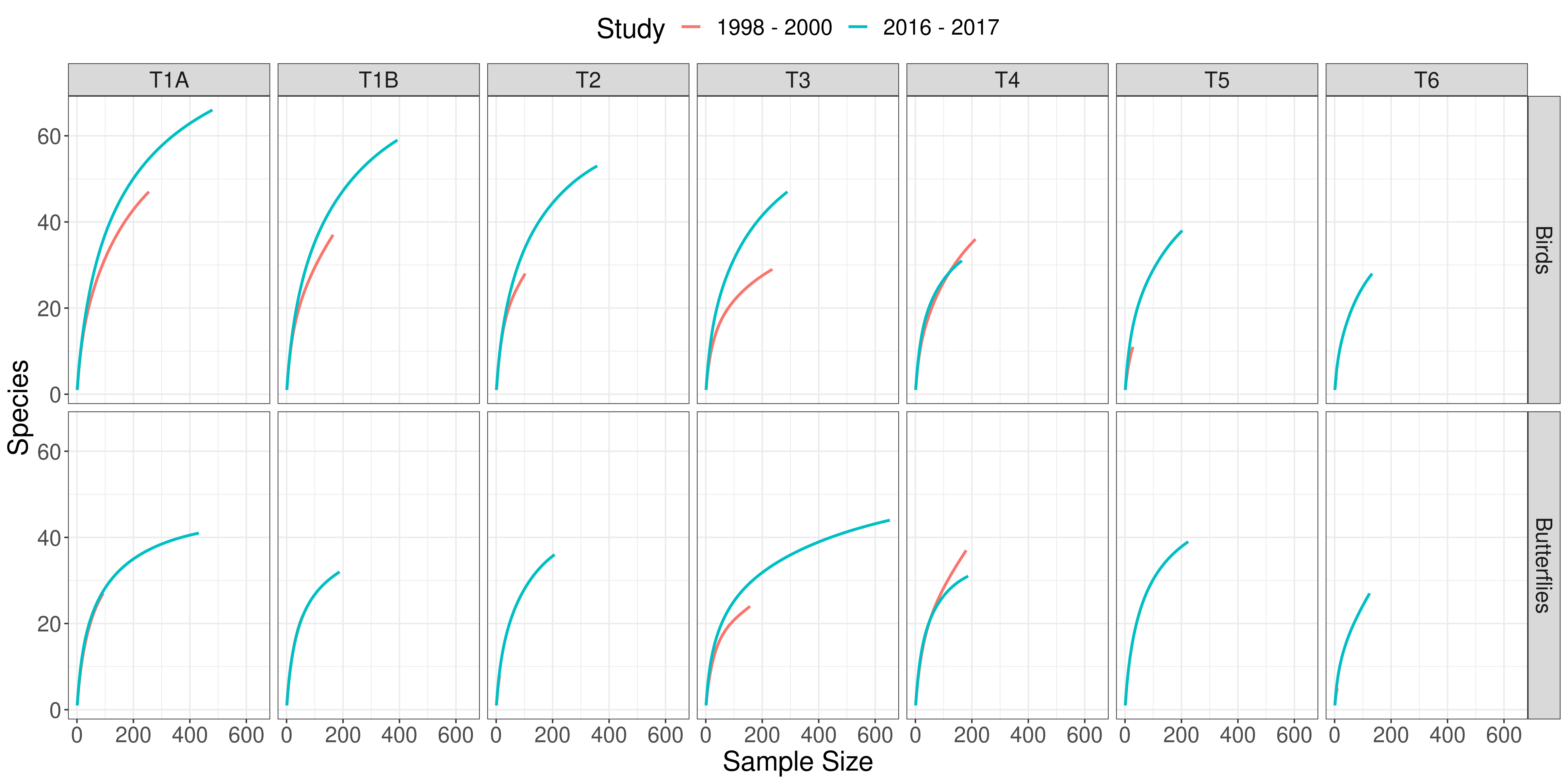

Figure A: Rarefaction curves indicating differences in sampling effort across studies by site.
